# Dog10K_Boxer_Tasha_1.0: A long-read assembly of the dog reference genome

**DOI:** 10.1101/2021.05.05.442772

**Authors:** Vidhya Jagannathan, Christophe Hitte, Jeffrey M. Kidd, Patrick Masterson, Terence D. Murphy, Sarah Emery, Brian Davis, Reuben M. Buckley, Yanhu Liu, Xiangquan Zhang, Tosso Leeb, Ya-ping Zhang, Elaine A. Ostrander, Guo-dong Wang

## Abstract

The domestic dog has evolved to be an important biomedical model for studies regarding the genetic basis of disease, morphology and behavior. Genetic studies in the dog have relied on a draft reference genome of a purebred female boxer dog named “Tasha” initially published in 2005. Derived from a Sanger whole genome shotgun sequencing approach coupled with limited clone-based sequencing, the initial assembly and subsequent updates have served as the predominant resource for canine genetics for 15 years. While the initial assembly produced a good quality draft, as with all assemblies produced at the time it contained gaps, assembly errors and missing sequences, particularly in GC-rich regions, which are found at many promoters and in the first exons of protein coding genes. Here we present Dog10K_Boxer_Tasha_1.0, an improved chromosome-level highly contiguous genome assembly of Tasha created with long-read technologies, that increases sequence contiguity >100-fold, closes >23,000 gaps of the Canfam3.1 reference assembly and improves gene annotation by identifying >1200 new protein-coding transcripts. The assembly and annotation are available at NCBI under the accession GCF_000002285.5.

## 1. Introduction

High quality reference genomes are fundamental assets for the study of genetic variation in any species. The ability to link genotype to phenotype and the subsequent identification of functional variants relies on high fidelity assessment of variants throughout the genome. This reliance is well illustrated by the domestic dog, which offers specific challenges for any genetic study. Featuring over 350 pure breeding populations, each breed is a mosaic of ancient and modern variants, and each reflects a complex history linking it to other related breeds. As a result of bottlenecks associated with domestication (15,000-30,000 years before present) and more recent individual breed formation (50 – 250 years before present) dog genomes contain long and frequent stretches of linkage disequilibrium (LD). While helpful for identifying loci of interest, long LD makes the necessary fine mapping for moving from marker to gene to variant both labor intensive and error prone.

In 2005, the first high-quality draft (7.5×) sequence of a Boxer dog, named Tasha, was made publicly available [1]. The reference sequence has proven useful in discoveries of canine associated molecular variants [2,3], including single-nucleotide variants (SNVs) and small indels, regulatory sequences [4,5], large rearrangements and copy number variants [6][7] associated with both inter and intra gene variation. The resulting SNV arrays, designed based on variation relative to the Tasha-derived assembly, have led to the success of hundreds of genome wide association studies (GWAS), advancing the dog as a system for studies of disease susceptibility and molecular pathomechanisms, evolution, and behaviour. However, for the dog system to advance further, a long read high quality assembly of a reference genome is needed. This will greatly improve the sensitivity of variant detection, especially for large structural variation. Furthermore, a high-quality assembly is an essential pre-requisite for accurate annotation, which is required to assay the potential functional effects of detected variants. Using the same dog as used for the initial assembly offers specific advantages, including the ability to integrate new findings with previous observations. A high quality genome assembly from the boxer Tasha will mean that the value of existing resources, such as existing bacterial artificial chromosome (BAC) libraries, and the wealth of experience and knowledge gained using previous versions of this dog’s genome, will be preserved for future research efforts. The dog genome assembly reported here was built using a combination of Pacific Biosciences (PacBio) continuous long read (CLR) sequencing technology, 10x Chromium linked reads, BAC pair-end sequences and the draft reference genome sequence CanFam3.1.

## 2. Materials and Methods

### 2.1. Whole Genome Sequencing

A single blood draw from which genomic DNA was isolated from blood leukocytes of a female Boxer, Tasha, which also was used to generate the previous CanFam 1, CanFam 2 and CanFam 3 genome assemblies was utilized here. Continuous long-read (CLR) sequencing was carried out at Novogene Bioinformatics Technology Co., Ltd (Beijing, China) with a PacBio Sequel sequencer (Pacific Biosciences, Menlo Park, CA, USA). Approximately 100 µg of genomic DNA were used for sequencing. SMRTbell libraries were prepared using a DNA Template Prep Kit 1.0 (PacBio), and 56 20-kb SMRTbell libraries were constructed. A total of 252 Gb of sequence data was collected. High molecular weight DNA from Tasha was also sequenced with Chromium libraries (10x Genomics, Pleasanton, CA, USA) on Illumina (San Diego, CA, USA) HiSeq X (2×150 bp), generating 589,824,390 read pairs or 176 Gb of data.

### 2.2. Genome Assembly Workflow

We assembled the genome using the Canu (v1.6) [8] and wtdbg2 [9] assembly algorithms. Briefly, the pipeline was composed of assembly, scaffolding and a final polishing step. PacBio reads had a mean read length of 8.5 kb and were used for the *de novo* assembly. The reads were corrected using the Canu error correction module which generates a consensus sequence for each read using its best set of long read overlaps. The corrected consensus reads were then assembled using the wtdbg2 algorithm[9], which is designed for assembly of long reads produced by the PacBio or Nanopore technologies. The assembled contigs were polished with raw PacBio reads using the WTPOA-CNS tool of the WTDBG2 package. This was followed by misassembly detection and correction with TIGMINT [10]. End sequences from BAC clones were extracted from the TraceDB of NCBI and used for scaffolding corrected contigs using the BESST algorithm (v2.2.8) [11]. Gap filling was done using the PacBio subreads with PBjelly (from PBSuite v15.8.24) [12] and one additional round of genome polishing was carried out using Pilon v1.23. [13] with the 10x Chromium reads. Finally, RaGOO (v1.1) [14] was used for reference guided scaffolding using CanFam3.1 as the reference. The draft scaffolds were subjected to additional gap closure using PBJelly.

### 2.3. Assembly Quality Control

The scaffold order and orientation of the assembly was assessed by aligning it to an existing radiation hybrid map (RH-map) comprising 10,000 markers [15]. A chromosome-wide review of scaffold discrepancies was determined visually, and those that were incorrectly ordered were corrected. The assembly was also assessed for completeness using BUSCO [34] which provides a summary of genome completeness using a database of expected gene content based on near-universal single-copy orthologs from mammalian species with genomic sequence data. This includes 4,104 single copy genes that are evolutionarily conserved between mammals.

#### 2.3.1 Fosmid end sequence alignment

End sequences from previously constructed fosmid libraries from Tasha were aligned to the assembly as previously described [1]. Concordant clones were considered to be those with an inward read orientation and a size between 35,328 and 43,453 bp. Using bedtools [16], the physical coverage of concordant clones in 5 kb windows along the genome was determined. Segments of the primary chromosome assemblies that were not supported by any concordant fosmids were also identified. Analysis was limited to the primary chromosome assemblies (chr1-chr38, chrX) and any interval that intersected with chromosome ends was discarded. This resulted in a total of 1004 regions, of which 282 intersected with a segmental duplication interval (considering the union of assembly and read-depth based annotations). To assess the significance of the intersection with segmental duplications, we performed 1,000 random permutations of the intervals using bedtools and found that 49 to 103 of the intervals intersected with a duplication, with a mean intersection rate of 75.

#### 2.3.2 Alignment of finished BAC clone sequences

A list of assembled BAC clones from the CH-82 library was obtained from ftp://ftp.ncbi.nih.gov/repository/clone/reports/Canis_familiaris/CH82.clone_acstate_9615.out. The sequence of 395 finished clones was aligned to the long read Tasha assembly using minimap2(v2.17)[17]. One clone (AC190394.3) did not have a minimap2 alignment, 124 clones returned mulitple alignment positions, 124 clones aligned to a single position annotated as duplicated in the Tasha4 assembly, and four clones returned alignments that did not include the entire BAC sequence. We therefore focused on a set of 142 clones that had alignment to a single locus based on minimap2 with a query alignment that encompassed the entire clone length and that did not overlap with regions annotated as segmental duplications in the Tasha4 assembly. An optimal global sequence alignment between the BAC sequence and the assembly was then determined using stretcher [18] with default parameters.

### 2.4. Detection of Common Repeats and Segmental Duplications

Common repeats were identified with RepeatMasker (v4.0.7) using the rmblastn (v2.2.27+) search engine and a combined repeat database consisting of the Dfam_Consensus-20170127 [19] and Rep-Base-20170127 [20] releases.

Segmental duplications in the assembly were detected using two approaches. First, duplicated regions were identified based on assembly self-alignment using the program SEDEF [21]. Duplications with a least 90% sequence identity and length of 1 kb were retained. Second, duplications were defined based on an analysis of the depth of coverage of Illumina sequencing data using the fastCN [22] program. Copy-number was estimated in non-overlapping windows each containing 3 kbp of unmasked sequence.

Control regions for normalization were converted to Dog10K_Boxer_Tasha_1.0 coordinates using the lift-Over tool [23,24]. Segmental duplications were defined as segments of four or more consecutive windows with an estimated copy-number of at least 2.5. Comparable annotations for the CanFam3.1 assembly were obtained from [25].

### 2.5. Gene Annotation

The assembly was annotated using the previously described NCBI pipeline [26][27]. The pipeline uses a WindowMasker-masked genome for building gene models substantiated with RNA-seq data and protein alignments. RNA-sequencing data from various dog tissues were used for the gene prediction (https://www.ncbi.nlm.nih.gov/genome/annotation_euk/Canis_lupus_familiaris/106/).

### 2.6 Genome assembly alignment

The Dog10K_Boxer_Tasha_1.0 assembly was aligned to the CanFam3.1 assembly using minimap2 (v2.17)[17] with the ‘asm5’ option. Insertions and deletions were identified using the paftools.js program distributed with minimap2 with default options. Analysis was restricted to the primary chromosome sequences (chr1-38 and chrX). Regions that overlapped with assembly gaps, segmental duplications detected based on assembly self-alignment, or segmental duplications identified by read depth were removed.

### 2.7 Structural variant detection

Raw PacBio reads were aligned to the CanFam3.1 and Dog10K_Boxer_Tasha_1.0 assemblies using minimap2 (v2.17)[17]. Structural variants were identified using sniffles (v1.0.12) [28]. Only calls with precise breakpoints on the primary chromosome sequences (chr1-38 and chrX) were considered. Calls were filtered to remove insertions and deletions that intersect with assembly gaps.

### 2.7 BAC assembly

Bacterial artificial chromosome (BAC) clones that mapped to the amylase locus were received from BACPAC resources center (Emeryville, CA). BACs were streaked to obtain single clones on LB agar with 100 ug/ul chloramphenicol and singe clones were cultured 20-24 hours at 37°C in 100ml LB broth with 100 ug/ul chloramphenicol. BAC DNA was isolated using NucleoBond Xtra Midi kit for transfection-grade plasmid DNA without NucleoBond® Finalizer (Machery-Nagel, Bethlehem, PA) and, after precipitation and drying, resuspended in 500 ul H_2_O by incubating 72-96 hours at 4°C. Within 48 hours of resuspension, BAC DNA was sequenced on a Minion with the Flongle adapter (Oxford Nanopore Technologies, Oxford, UK). Libraries were made using the Rapid Barcoding Sequencing kit (Oxford Nanopore Technologies, SQK-RBK004) according to manufacturer’s protocol, except, for fragmentation where 0.25 ul of Fragmentation Mix was mixed with 200 ng of DNA in 4.75 ul of water, incubated 30° C for 1 minute then 80° C for 1 minute, and cooled on ice. Following fragmentation, BAC libraries were pooled by adding 1.67 ul of each library prep, 0.5 ul Rapid Primer (RAP) was added, and the mix was incubated for 5 minutes at room temperature. Flow cells were primed and loaded according to manufacturer’s protocol.

Nanopore reads from the BAC were assembled using the pipeline described in https://github.com/KiddLab/run_canu_bac. Briefly, raw reads were filtered for hits to *E. coli* and assembled using canu (v2.1)[8]. The unique portion of the resulting circular contig was then extracted and polished using racon (v1.4.10)[29]. Finally, the vector backbone sequence was removed and the contig was rotated to begin at the appropriate position. The final CH82-451P03 sequence was compared to the Dog10K_Boxer_Tasha_1.0 assembly using MUMmer (v3.23)[30].

### 2.8. Mapping SNV Array Probes

Chromosomal sequences from CanFam3.1 and Dog10K_Boxer_Tasha_1.0 were aligned to each other using blat [31]. The aligned fragments were processed using UCSC tools to create the necessary chain file for use with the liftOver tool. The liftOver was performed using the default settings with the “-multiple” option included. Genomic positions from both the Affymetrix (Santa Clara, CA, USA) Axiom Canine HD Array and Illumina (San Diego, CA, USA) CanineHD BeadChip were converted from CanFam3.1 to Dog10K_Boxer_Tasha_1.0. Genomic positions were obtained for the Axiom Canine HD Array from: https://sec-assets.thermofisher.com/TFS-Assets/LSG/Support-Files/Axiom_K9_HD.na35.r5.a7.annot.csv.zip; and for the CanineHD Bead-Chip:ftp://webdata2:webdata2@ussd-ftp.illumina.com/downloads/ProductFiles/CanineHD/CanineHD_B.csv. The bed files resulting from the lift over were converted to Plink map files. All markers were included in each map file and markers were ordered sequentially according to the order they were downloaded from their corresponding URLs. Markers for which no position was obtained were placed on chromosome “0” at position “0”.

## 3. Results

DNA isolated and stored at -80°C at NHGRI from the same female Boxer, Tasha, used for the CanFam 3.1 draft genome sequence was utilized to generate a new assembly. Frozen DNA from the same aliquot was thawed and used to prepare high molecular weight DNA libraries, which were sequenced using PacBio single-molecule real-time (SMRT) and 10x Genomics Linked-Reads sequencing technologies. Approximately 100-fold coverage (252 Gb) and 74-fold coverage (176 Gb) of the genome were generated using PacBio and 10x Genomics reads, respectively.

### 3.1. Dog10K_Boxer_Tasha_1.0 assembly

PacBio SMRT cells produced 27,878,642 reads with a mean length of 8,514 bp and N50 read length, length at which 50% of the bases are in reads longer or equal to, was 13,189 bp. All PacBio reads were used for the assembly. The assembly pipeline (Fig. 1) underwent initial read correction with Canu. After correction, 5,586,195 reads were used for assembling with wtdbg2, obtaining a corrected read cut-off of 14 kb that provided 43-fold (104,569,563,638 bases) genome coverage for input. The initial ungapped assembly of WTDBG2 contained 1,562 contigs with an N50 of 23.8 Mb. Tigmint (v.0.4) was used to correct initial assembly errors by incorporating the linked reads generated by 10x Genomics Chromium long read technology. Tigmint split 75 missambled contigs, which resulted in an assembly featuring 1786 contigs, of which 1724 were >500 bp. The assembly contig N50, the contig length in the assembly that equal or longer contigs contain half the bases of the genome, was 23.72 Mb.

**Figure 1.**
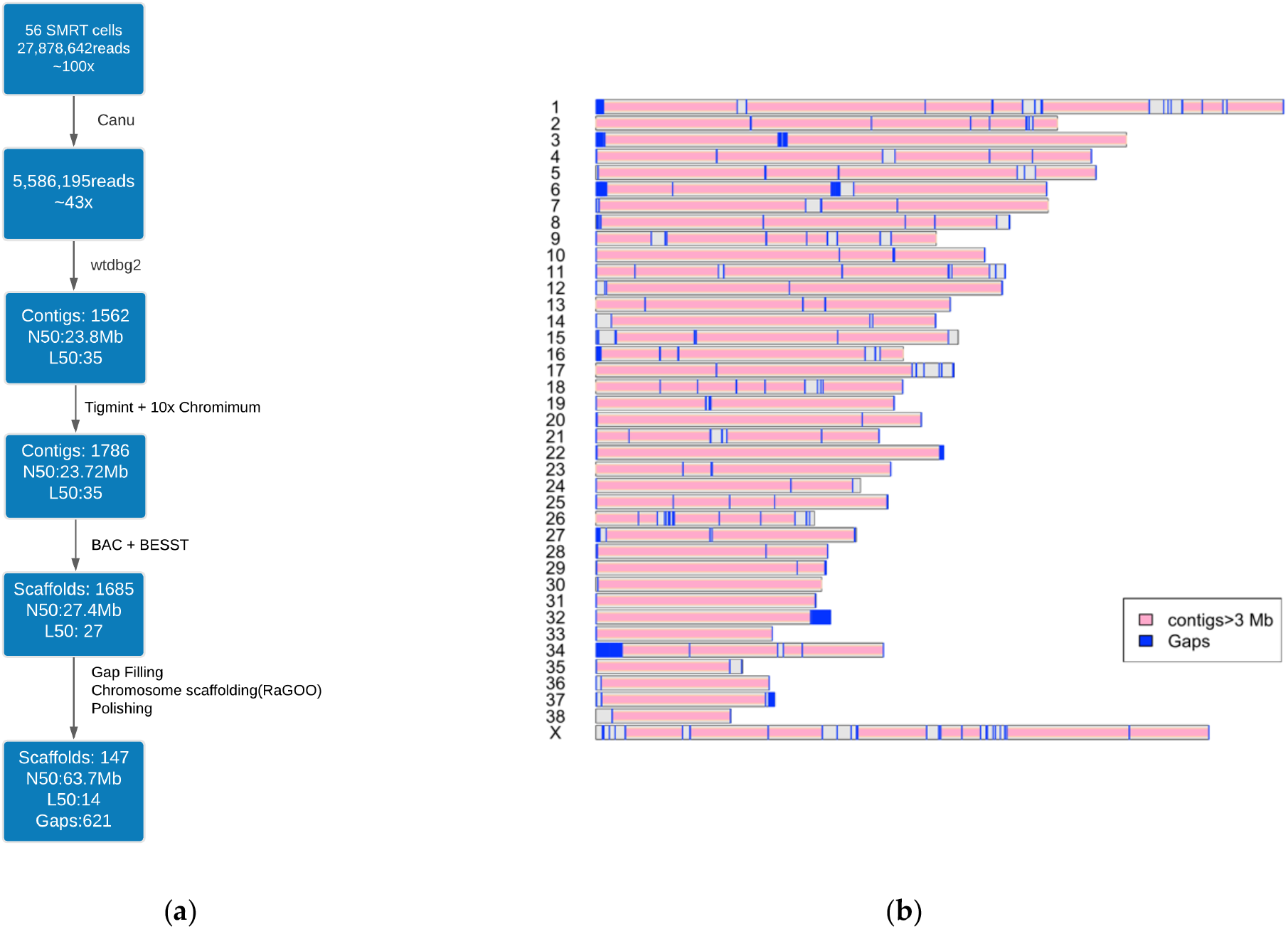
Dog_10k_Boxer_Tasha_1.0 assembly. (**a**) Assembly pipeline (**b**) Ideogram showing chromosomes, contigs, gaps.

The Tigmint corrected assembly was then scaffolded with BAC end sequences. The resultant scaffolding, constructed with the BESST algorithm (v 2.2.8) resulted in an assembly of 1685 scaffolds, which increased the N50 to 27.4 Mb. RaGOO was then used to scaffold the data into 39 chromosomes based on CanFam3.1. The chromosome level scaffolds had a minimum of four contigs as noted on chromosomes 28, 30 and 36 and a maximum of 82 contigs on the X chromosome. The N50 of the scaffolded assembly was 63,738,581 bp (Table 1). The assembly contained 621 spanned gaps closing >23,000 of the Canfam3.1 assembly (18.25 Mb) (Fig. 1).

**Table 1.** Summary statistics for the Dog10K_Boxer_Tasha_1.0 genome assembly and comparison with current dog reference genome CanFam3.1.

| **Statistic** | **CamFam3.1** | **Dog10K_Boxer_Tasha_1.0** |
| --- | --- | --- |
| Total sequence length | 2,410,976,875 | 2,312,802,206 |
| Total ungapped length | 2,392,715,236 | 2,312,743,367 |
| No. of scaffolds | 3,310 | 147 |
| No. of unplaced scaffolds | 3,228 | 107 |
| Scaffold N50 | 45,876,610 | 63,738,581 |
| Scaffold L50 | 20 | 14 |
| No. of unspanned gaps | 80 | 399 |
| No. of spanned gaps | 23,796 | 621 |
| No. of contigs | 27,106 | 1162 |
| Contig N50 | 267,478 | 27,487,084 |
| Contig L50 | 2,436 | 31 |
| No. of chromosomes* | 39 | 39 |

The quality of the Dog10K_Boxer_Tasha_1.0 assembly was assessed by comparison with an existing RH-map of 10,000 markers. The comparison strongly supported the overall accuracy of the assembly. There were two major discordances between the RH map and the draft assembly order of the contigs, one on chromosome 6 and the other on chromosome 11. The order was corrected, and gaps were again closed using PBJelly and PacBio SMRT raw reads. Discrepancies involving blocks of ∼1 Mb on chromosome 9 and 0.2 Mb on chromosome 16 could not be resolved and will require further investigation.

### 3.2 Assembly quality assessment

We used fosmid clone end sequences to identify regions that may be misassembled in Dog_10k_Boxer_Tasha_1.0. We identified 895,746 clones with a concordant mapping based on the orientation of the end-sequences and the apparent size of the cloned fragment (Figure S1), yielding a median genomic physical coverage of 17 concordant clones (Figure S2). Using this map of fosmid coverage, we identified 1,004 intervals (32.5 Mb) on the primary chromosomes that do not intersect with a concordantly mapping fosmid (Table S1). We found that 282 of these intervals intersected with regions of segmental duplication in Tasha4, a value greater than that observed in any of 1,000 random permutations. This indicates that duplicated regions are enriched for potential missassembly.

We also assessed the per-bp sequence accuracy of the Dog_10k_Boxer_Tasha_1.0 assembly using 142 finished BAC clones from the CH-82 library that have a unique alignment to Dog_10k_Boxer_Tasha_1.0. Discarding alignment gaps, mismatches were observed at 14,255 of 26,778,153 aligned nucleotides (Figure S3). Assuming that all mismatches represent errors in the Dog_10k_Boxer_Tasha_1.0 sequence, a conservative assumption since heterozygous sites as well as errors in the BAC sequence are expected, the observed mismatch rate corresponds to an estimated per-base sequence quality [32] of Q33. We note, however, that the apparent number of alignment gaps is higher than the apparent single base substation rate, suggesting that indels remain the primary error mode in long-read assemblies (Table S2).

### 3.3 Assembly Completeness

The completeness of the assembly was assessed using BUSCO, which uses a set of universal single-copy orthologs. This analysis showed an improvement of BUSCO completeness from 92.2% in CanFam3.1 to 95.3% in Dog10K_Boxer_Tasha_1.0 (Table 2).

**Table 2.** Comparison of BUSCO analysis of genomes

| Statistic | Dog10k_Boxer_Tasha_1.0 | CanFam3.1 |
| --- | --- | --- |
| Complete BUSCOs | 95.3% | 92.2% |
| Complete and single copy BUSCOs | 94.1% | 91.1% |
| Complete and duplicated BUSCOs | 1.2% | 1.1% |
| Fragmented BUSCOs | 2.1% | 4.0% |
| Missing BUSCOs | 2.6% | 3.8% |

We further compared the structural accuracy of the RaGOO arranged chromosome-level scaffolds to that of the CanFam3.1 chromosomes. We identified several regions known to be mis-assembled in CanFam3.1 and were now corrected in the Dog10k_Boxer_Tasha_1.0 assembly. These regions were supported by corresponding BAC end sequences (Figure S4).

Additionally, the orientation of chromosomes 27 and 32 were reversed compared to CanFam3.1. The two chromosomal re-orientations were backed by evidence in [33] and [34] based on recombination rates in dog chromosomes and fluorescence *in situ* hybridization experiments by Matthew Breen (personal communication).

### 3.4. Gene Annotation

Annotation of the Dog10K_Boxer_Tasha_1.0 assembly was carried out using the NCBI annotation pipeline and released via the NCBI ftp site [35]. The annotation pipeline used RNA-seq data from more than 25 tissues, along with known RefSeq, Genbank transcripts and canine expressed sequence tags. Statistics from the annotation release 106 are listed in Table 3. The annotation includes 20,100 protein coding genes, which is comparable to annotations of other carnivores (average 20,105, stdev 1078, from 27 species). A total of 1299 protein-coding transcripts from 737 genes were identified as novel as they do not align to CanFam3.1 assembly. We found 78 out of 2,473 known RefSeq transcripts did not map to the Dog10k_Boxer_Tasha_1.0 assembly[35]. Significantly, we observed a 7.0% increase (17,721 vs 16,554) in the number of annotated protein-coding genes with very high coverage (>=90%) alignments compared to their best hits in SwissProt, with 88% of all protein-coding genes having at least one isoform exceeding 90% coverage. In addition, the new Tasha assembly has only 4.5% (891) of protein-coding genes represented with corrected models that compensate for suspected frameshifts or premature stop codons in the genome, compared to 5.5% for the prior NCBI annotation of CanFam3.1, or 5.6 - 11.3% for NCBI annotations of several other canine assemblies. These improvements can be largely attributed to fewer assembly gaps and the fact that gaps comprising exons of several genes have now been closed (Figure S5). For example, 5770 genes in CanFam3.1 have gaps within and flanking them. Only 12 of these genes still have gaps overlapping their exons and introns in Dog_10k_Boxer_Tasha_1.0.

**Table 3.** Annotation statistics for NCBI annotation release 106. * are non-coding RNA genes that cannot be classified.

| Feature | Dog10k_Boxer_Tasha_1.0 /<br>annotation release 106 |
| --- | --- |
| Protein coding genes | 20,100 |
| Non-coding genes | 15,306 |
| Small non-coding genes | 2,083 |
| Long non-coding genes | 12,667 |
| Miscellaneous* non-coding genes | 10 |
| Pseudogenes | 4887 |

### 3.5 SNV Array Probes mapped to Dog10k_Boxer_Tasha_1.0

Marker positions from the Axiom Canine HD Array and CanineHD BeadChip were mapped from CanFam3.1 to Dog10K_Boxer_Tasha_1.0. For the Axiom Canine HD Array and CanineHD BeadChip, 98.12% and 97.91% of markers, respectively, were successfully mapped to the new assembly. The data is available as supplementary files S1 and S2. The majority of markers on both arrays mapped to the same chromosome on both assemblies, with marker order remaining mostly intact. The largest contiguous off-diagonal collection of markers was found on chromosome 16 in CanFam3.1 and on chromosome 34 in Dog10K_Boxer_Tasha_1.0.

### 3.6. Analysis of duplications

We identified segmental duplications in the Dog10K_Boxer_Tasha_1.0 assembly using two approaches. First, based on assembly self-alignment, we defined segmental duplications as segments at least 1 kb in length with a sequence identity of 90% or greater. This identified 5,730 intervals encompassing 28.7 Mb of sequence on the primary chromosome assemblies (Table S3). Second, we identified 321 intervals encompassing 38.3 Mb of sequence based on excess depth of coverage from Illumina sequencing reads. Both of these measures of duplication content are less than found in the Great Dane Zoey or CanFam3.1 assemblies [25], indicating that these duplicated sequences are not correctly resolved in the Dog10K_Boxer_Tasha_1.0 genome assembly.

### 3.7 Analysis of repetitive sequences

We identified common repeats in the Dog10K_Boxer_Tasha_1.0 assembly using RepeatMasker. A total of 41.1% of the assembly is comprised of repeats, with most falling into one of three categories: LINEs (469 Mb), SINEs (241 Mb) and LTRs (110 Mb). A complete summary of the repeat element composition is available in Table 4. We compared the results with an equivalent annotation of CanFam3.1. As before, we limited analyses to the primary chromosome sequences. At a high level, the repeat content of the Dog10K_Boxer_Tasha_1.0 and CanFam3.1 assemblies is similar (Table 4). However, the primary chromosome sequences in the Dog10K_Boxer_Tasha_1.0 assembly includes substantially more sequence classified as ‘satellite’, reflecting the ability of long-read sequencing to extend into subtelomeric and pericentromeric chromosomal regions. Unexpectedly, RepeatMasker analysis indicated that CanFam3.1 contains more sequence annotated as short interspersed nuclear elements (SINEs) while Dog10K_Boxer_Tasha_1.0 contains more sequence annotated as long interspersed nuclear element (LINEs). Sequences belonging to these repeat types make up a substantial fraction of the canine genome. LINE and SINE retrotransposons move via a copy-and-paste mechanism and new insertions accumulate mutations over evolutionary time scales [36]. Focusing on the youngest sequences shows that CanFam3.1 contains over 9,000 more copies of a family of carnivore SINEs (SINECs) that show less than 10% sequence divergence, while the Dog10K_Boxer_Tasha_1.0 assembly contains 576 more LINEs that have less than 10% sequence divergence and are longer than 4 kb. We aligned the Dog10K_Boxer_Tasha_1.0 and CanFam3.1 assemblies to further explore this difference in SINE and LINE content and identified 55,329 insertion-deletion differences between the assemblies longer than 10 bp. The variant size distribution has clear peaks corresponding to the expected sizes of dimorphic LINEs and SINEs (Figure 2).

**Table 4.** Repeat content of the Dog10K_Boxer_Tasha_1.0 and CanFam3.1 assemblies. Results are shown for the primary chromosome sequences.

|  | Dog10K_Boxer_Tasha_1.0 |  | CanFam3.1 |  |
| --- | --- | --- | --- | --- |
| Repeat Class | Elements | bp | Elements | bp |
| DNA | 341,866 | 65,043,282 | 347,025 | 65,997,048 |
| LINE | 1,286,663 | 467,394,285 | 1,307,498 | 470,518,469 |
| LTR | 378,505 | 111,520,139 | 384,551 | 113,151,392 |
| Low_complexity | 123,075 | 6,525,287 | 120,803 | 6,009,804 |
| RC | 1,636 | 345,889 | 1,649 | 347,342 |
| RNA | 489 | 103,097 | 504 | 105,770 |
| SINE | 1,579,792 | 240,791,186 | 1,605,511 | 244,461,861 |
| Satellite | 5,730 | 11,298,647 | 635 | 624,881 |
| Simple_repeat | 891,331 | 40,450,974 | 895,091 | 38,358,719 |
| Unknown | 3,449 | 559,562 | 3,487 | 565,722 |
| rRNA | 953 | 129,078 | 958 | 115,711 |
| scRNA | 70 | 4,996 | 71 | 5,156 |
| snRNA | 4,492 | 278,022 | 4,617 | 285,578 |
| srpRNA | 45 | 8,900 | 47 | 9,496 |
| tRNA | 35,501 | 2,608,084 | 35,906 | 2,636,278 |

**Table 5.** Repeat content for the lowly diverged SINE and LINE sequences

|  | Dog10K_Boxer_Tasha_1.0 | CanFam3.1 |
| --- | --- | --- |

| Repeat Class | Elements | bp | Elements | bp |
| --- | --- | --- | --- | --- |
| SINEC | 1,125,416 | 177,104,238 | 1,146,663 | 180,147,553 |
| SINEC < 10% divergence | 454,869 | 71,490,885 | 464,113 | 72,819,234 |
| LINE/L1 | 853,212 | 379,452,954 | 869,259 | 381,738,114 |
| LINE/L1 < 10% divergence<br>and >= 4kb | 4,805 | 26,935,018 | 4,229 | 23,359,516 |

**Figure 2.**
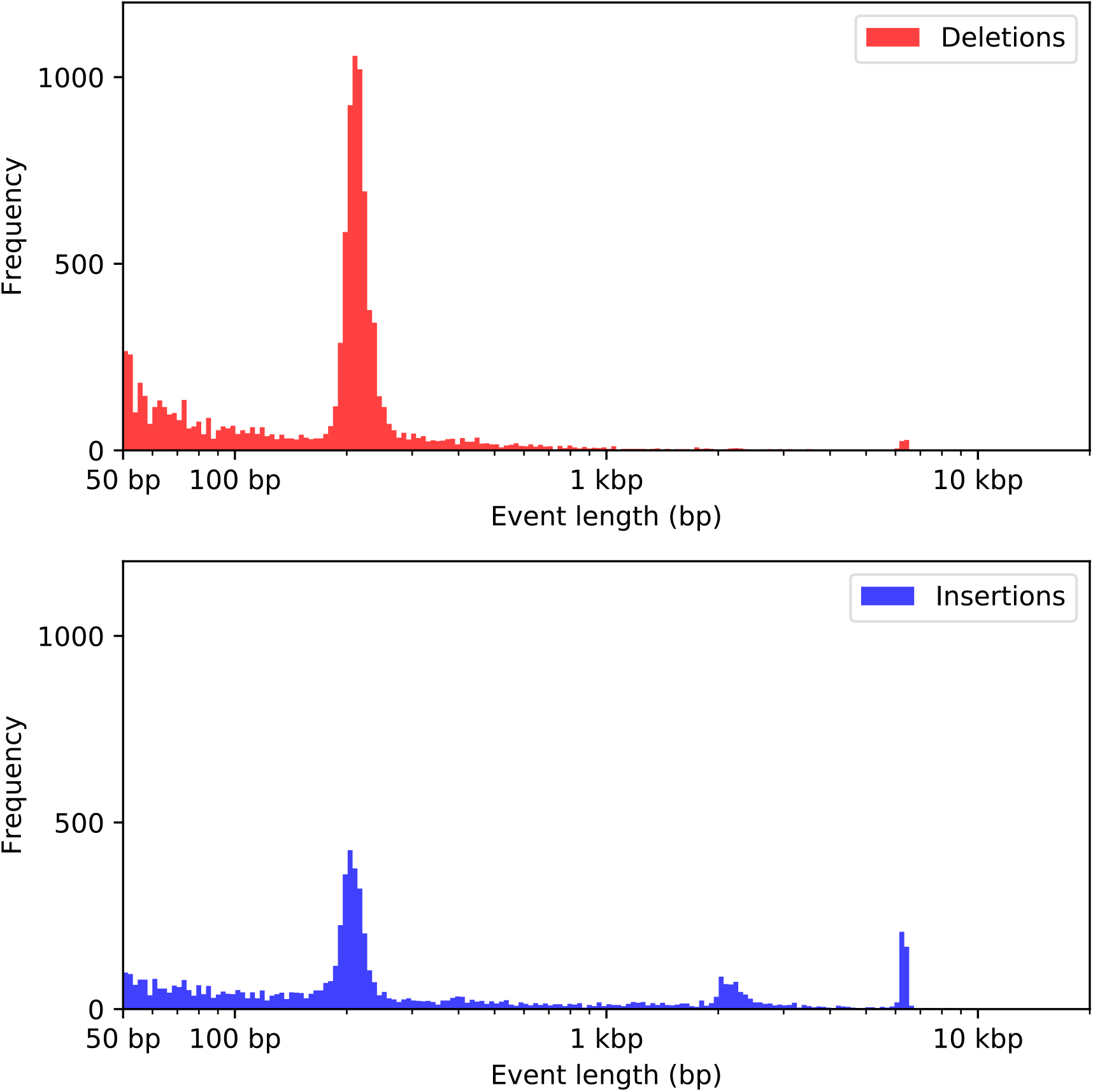
Size distribution of insertion-deletion differences identified between the Dog10K_Boxer_Tasha_1.0 and CanFam3.1 assemblies. The sizes of 22,330 sequence present in CanFam3.1 but absent in Dog10K_Boxer_Tasha_1.0 (red, deletions) and of 32,999 sequences present in Dog10K_Boxer_Tasha_1.0 but absent in CanFam3.1 (blue, insertions) are shown. The bins of each histogram are of equal size on a logarithmic scale.

Since LINEs and SINEs insertions are highly polymorphic among canines [25,37], we reasoned that the representation in the Dog10K_Boxer_Tasha_1.0 and CanFam3.1 assemblies may reflect the differential inclusion of heterozygous insertions. To assess this possibility, we identified structural variants relative to each assembly using the Tasha PacBio reads. Given the challenges associated with accurately discovering large insertions, we focused our analysis on deletion variants. We identified 35,187 deletions based on alignment to CanFam3.1 and 26,667 deletions based on alignment to Dog10K_Boxer_Tasha_1.0 (Supporting Files S3 and S4). Analysis of the variant size distribution is consistent with differential representation of heterozygous SINEs and LINEs in the two assemblies: there is an excess of ∼200 bp deletions when mapping to CanFam3.1 while there is an excess of ∼6 kb deletions when mapping to Dog10K_Boxer_Tasha_1.0 (Figure 3).

**Figure 3.**
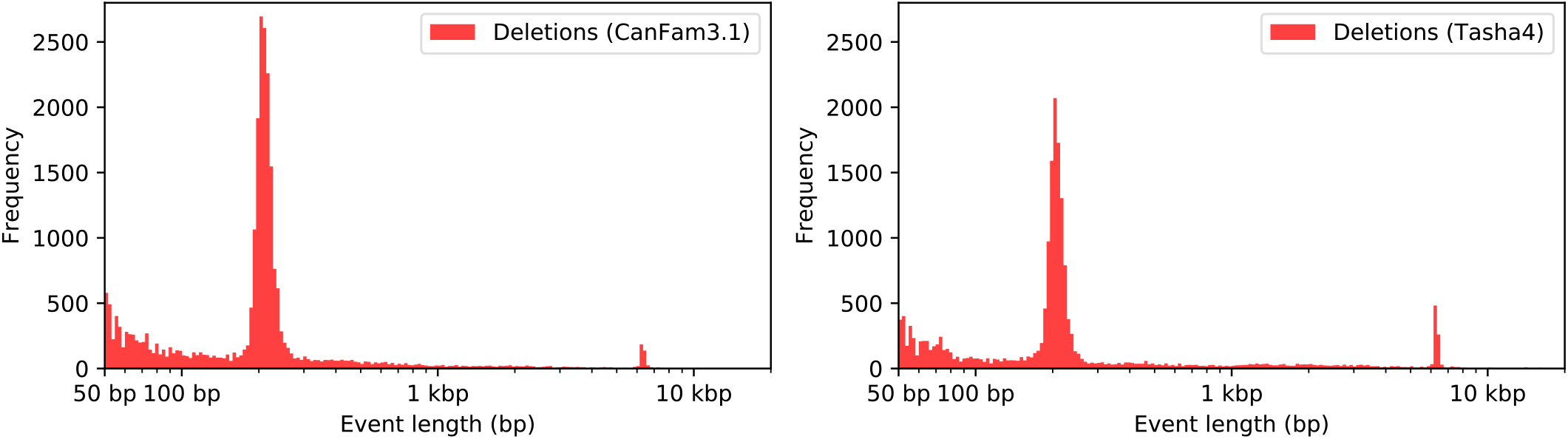
Discovery of deletion variants using PacBio reads. Deletions were identified based on alignment of PacBio reads to the CanFam3.1 (left) or Dog10K_Boxer_Tasha_1.0 (right) assemblies. The bins of each histogram are of equal size on a logarithmic scale.

### 3.8. Duplications at the Pancreatic Amylase Locus

Changes in amylase copy number and expression have been correlated with dietary preferences across mammals [38]. Increased copy number of the gene *AMY2B*, which encodes pancreatic amylase, has been associated with adaptation to a starch-rich diet in modern dogs *[39–41]. AMY2B* copy number is variable both within and among modern dog breeds [42], suggesting a dynamic copy number state, perhaps reflecting recurrent expansion and contraction of a tandemly duplicated array. Long-read assembly data from a Basenji, named China [43], and a German Shepherd, named Nala [44], support the presence of a tandemly duplicated architecture at the *AMY2B* locus. In addition to tandem duplications, large segmental duplications encompassing *AMY2B* have also been described [22,45].

In the Dog10K_Boxer_Tasha_1.0 assembly *AMY2B* is represented as a single copy on chromosome 6. Using Illumina read data, we estimate that the diploid *AMY2* copy number in Tasha is 12 (Figure 4). We found that Tasha is also heterozygous for a large duplication encompassing this locus. Examination of aligned fosmid end-sequence pairs revealed two clusters of clones that have an everted orientation consistent with a tandem duplication structure [46]. We identified the boundaries of these tandem duplications using the raw PacBio reads, defining the boundaries of tandem duplication units that are 1.9 Mb and 14.9 kb in length. Due to the presence of the larger duplication, the 12 *AMY2B* copies found in the Dog10K_Boxer_Tasha_1.0 genome are distributed among three structural alleles. Using Nanopore sequencing, we assembled a BAC mapping to this locus (CH82-451P03), and found that it contains a single copy of the *AMY2B* gene (Figure S6). Thus, at least one of the three structural *AMY2B* alleles in Tasha contains a single copy of this gene.

**Figure 4.**
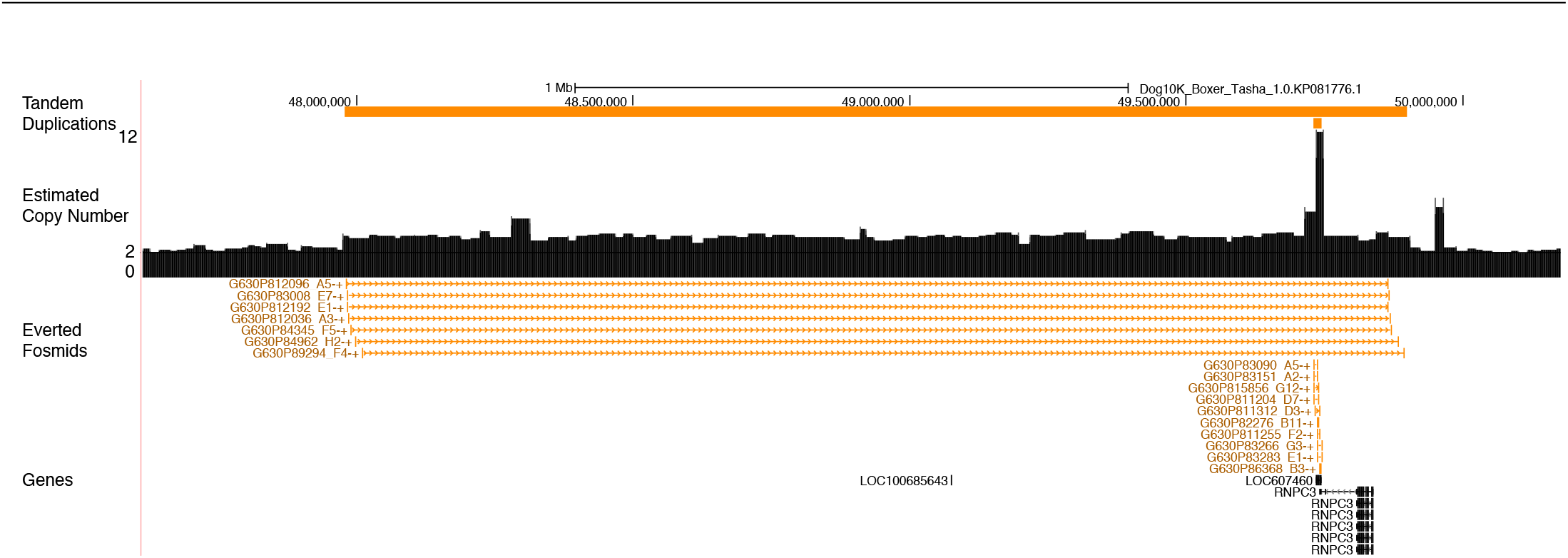
Structural variation at the amylase locus. A genome browser view illustrating structural variation at the amylase locus in Tasha is shown. The orange bars at the top indicate the locations of tandem duplications identified using the raw PacBio long-read data. This includes a large, 1.9 Mbp duplication (chr6:47977592-49898283) as well as a 14.8 kbp duplication (chr6:49729008-49743863). A read depth profile showing copy-number estimated from Illumina sequencing data is depicted as a bar plot across the interval. An elevated copy number of 3, corresponding to the 1.9 Mb duplication, is observed, as well as a spike in copy number overlapping with the *AMY2B* gene. Mappings of discordant fosmid end sequences are shown in orange below the copy number profile. Each depicted clone has end sequences that align in an everted orientation consistent with the presence of a tandem duplication. The position of gene models derived from the NCBI gene annotation, release 106, are shown at the bottom of the figure. The *LOC607460* gene model corresponds to pancreatic alpha-amylase (*AMY2B*).

## 4. Discussion

*Canis lupus familiaris*, the domestic dog, is now well-established as a genetic system for studies of disease susceptibility, physiology and morphology, all of which inform our understanding of human health. Major advances in human disease genetics have resulted directly from observations made in the dog. Some prominent examples include the identification of *PNPLA1* variants in human patients with autosomal recessive congenital ichthyosis 10 that was enabled by results obtained in Golden Retrievers [47] or the elucidation of the role of the *PRCD* gene in dogs with progressive cone-rod dystrophy and human patients with retinitis pigmentosa [48]. In addition, because of the availability of a canine genome assembly, canine disease models are now well established for several diseases including Duchenne muscular dystrophy [49], hypohidrotic ectodermal dysplasia[50], and Leber congenital amaurosis [50]. Similarly, canid evolution has revealed new insights as to shifts in canine behaviors that are both surprising and informative, and evinced human dependence on dogs for early survival. While early canine studies relied on segregation studies in families, and later GWAS studies in case control cohorts, the most informative studies now rely on large numbers of SNVs and small indels retrieved from publicly available sequences aligned to the reference genome. As such, the reference genome is of critical importance, as current sequence-based GWAS studies highlight not just gene regions, but genic or regulatory variants of interest.

Using PacBio and 10x Chromium long-reads, Dog10k_Boxer_Tasha_1.0 was generated as a new dog genome resource, with a dramatically increased continuity. CanFam3.1 had a contig size of only 267 kb while the Dog10k_Boxer_Tasha_1.0 assembly has an N50 contig size of 27.3 Mb featuring a >100-fold increase in sequence continuity. The improvements in the Dog10k_Boxer_Tasha_1.0 genome sequence relative to the CanFam3.1 assembly included not only greater continuity and fewer gaps, but also led to the correction of misassembled gene regions like *OCA2* (Fig S4) which were supported by concordant alignments of BAC end sequences to the Dog10k_Boxer_Tasha_1.0 assembly.

The improvements in continuity and quality yielded a stronger template for annotation, resulting in better gene models. There is a 7.0% increase in protein-coding genes with high-coverage (>=90%) alignments to SwissProt, likely resulting from the increased contiguity, and the percentage of protein-coding genes annotated with corrections for suspected frameshifts or premature stop codons is the lowest of any current canine assembly (4.5%, vs 5.6 – 11.3%), which may reflect the use of CLR reads and an additional polishing step. There are 78 of 2,743 known RefSeq transcripts (2.8%) that do not map to Dog10k_Boxer_Tasha_1.0 assembly, which is higher than observed for other assemblies and requires further investigation. In particular, whole genome alignments between Dog10k_Boxer_Tasha_1.0 and previous Tasha assemblies highlight two major deletions on the X chromosome in the new assembly: an 8 Mb deletion (NC_051843.1: 14.2M..22.2M) and a 4.5 Mb deletion (NC_051843.1: 72M..76.5M). Additional sequencing of the X chromosome is required to resolve these regions.

There is a systematic underrepresentation of GC-rich sequences in CanFam3.1, as the necessary cloning and sequencing steps did not amplify GC-rich DNA particularly well. Long-read sequencing for the new assembly did not use any cloning steps or PCR and, as a result, GC-rich sequences are better represented and many gaps that were present in CanFam3.1 could be closed. This is critical as GC-rich sequences are often found in the first exons and promoter regions of genes, and play important roles in regulation, such as through differential methylation of CpG islands. As a result, the Dog10k_Boxer_Tasha_1.0 assembly will allow for more accurate identification of genetic variation in GC-rich regulatory regions and methylome studies.

To date, five long-read based *de novo* dog genome assemblies [25,43,51] have been made available at the NCBI genome repository with comparable parameters such as number of genes annotated and number of gaps between the new assemblies. The NCBI has annotated all five genomes and made them available on their genome browser https://www.ncbi.nlm.nih.gov/genome/gdv/?org=canis-lupus-familiaris. The comparative results indicate a strong likelihood that more protein-coding transcripts, pseudogenes, and non-coding genes remain to be discovered and annotated. However, the highly continuous genome sequence reported here provides a greatly improved framework which will enhance characterization of functional sequences, genetic variation, and improve the utility of the thousands of canid sequences already generated, setting the stage for genetic studies of high accuracy and resolution.

Availability of *de novo* assemblies from different breeds will help to characterize structural variants (SVs), including the copy-number variations (CNV), mobile element diversity, chromosomal rearrangements, missing sequences and non-redundant sequences. In all species and, especially in dogs, a single reference genome from one individual is unable to represent the full spectrum of divergent sequences in populations worldwide. Dog genomes vary in both gene content, including tandem duplicated genes, CNVs distributed throughout the genome and in repetitive parts of the genome such as transposable elements. By characterizing genetic and structural variation within the canine species, *de novo* assemblies will better reveal the extensive variation in genome content among canine sub-populations defined by breeds, clades, and geography. The extensive analysis of the genetic variability of the canine genome will constitute the next paradigm shift for canine genomics.

## 5. Conclusions

We provide the Dog10k_Boxer_Tasha_1.0 genome assembly derived from the female boxer Tasha; the same dog that was used for the previous genome assemblies CanFam1, 2 and 3. Our assembly represents a sub-stantial improvement in continuity and completeness and, together with the associated annotation, will be a valuable resource for canine and comparative genetics research.

## Supplementary Materials

**Figure S1**. Apparent fosmid library insert size

**Figure S2**. Coverage of concordant fosmid clones

**Figure S3** Apparent sequence errors based on finished BAC clones

**Figure S4**. Corrected misassembled regions

**Figure S5** Filled Gaps in exons

**Figure S6** Alignment of CH82-451P03 to the Tasha4 assembly

**TableS1** Regions with no concordant fosmid coverage

**Table S2** BACalignedStats.xlsx

**TableS3** SegDup-align-table

**File S1** Axiom Canine HD Array map file for 10k_Boxer_Tasha

**File S2** CanineHD BeadChip map file for 10k_Boxer_Tasha

## Author Contributions

For research articles with several authors, a short paragraph specifying their individual contributions must be provided. The following statements should be used “Conceptualization, E.A.O, Y.Z and G.W; methodology, V.J, J.K. S.E, C.H, B.D; sequencing: G.W, S.E, J.K ;software, V.J and J.K.; formal analysis, V.J, J.K, C.H, Y.L and X.Z.; resources, E.A.O, Y.Z and G.W.; writing—original draft preparation, V.J T.L J.K and E.A.O.; writing—review and editing, V.J, T.L, J.K, E.A.O, G.W .; funding acquisition, E.A.O and Y.Z. All authors have read and agreed to the published version of the manuscript.” Please turn to the CRediT taxonomy for the term explanation. Authorship must be limited to those who have contributed substantially to the work reported.

## Data Availability Statement

The genome assembly is deposited at NCBI under accession number GCF_000002285.5. The associated BioProject accession number is PRJNA13179. BAC clone sequence has been deposited under accession MW972226.

