## Supplemental Figures for "Dog10K_Boxer_Tasha_1.0: A long-read assembly of the dog reference genome"

**Supporting information to Jagannathan et al.**

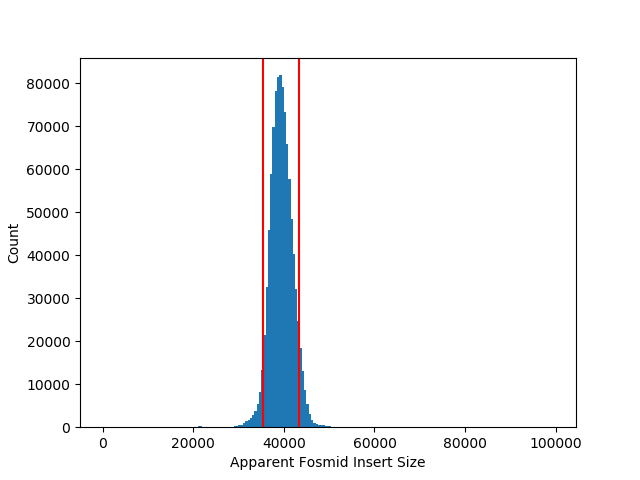

**Figure S1. Apparent fosmid library insert size**

Fosmid clone end-sequences were aligned to the primary assembly and the distribution of apparent clone insert size determined from clones with a proper “inward” mapping orientation. The median insert size was 39.39 kb with a median absolute deviation of 1,625 bp. We defined concordant clones as those with a proper read orientation and with an apparent size within 2.5 median absolute deviations of the median (35,328 – 43,453 bp, red lines). A total of 895,746 clones, representing 90.3% of those considered, were deemed to be concordant.

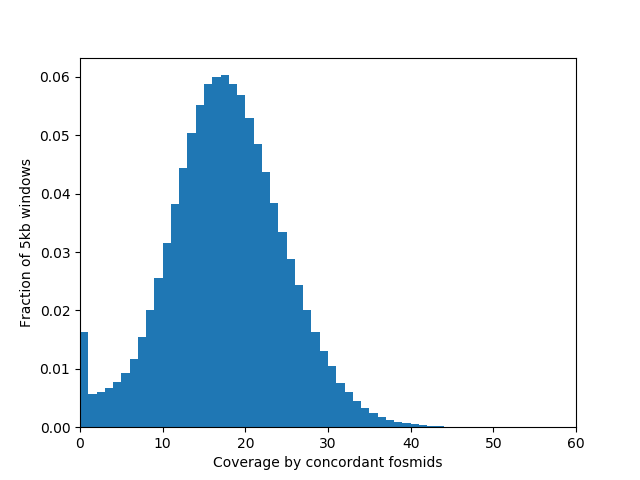

**Figure S2. Coverage of concordant fosmid clones**

The distribution of physical coverage of concordant clones across 5,000 bp genomic bins is shown. Only the primary chromosome sequence assemblies were considered. The median coverage level is 17 concordant clones.

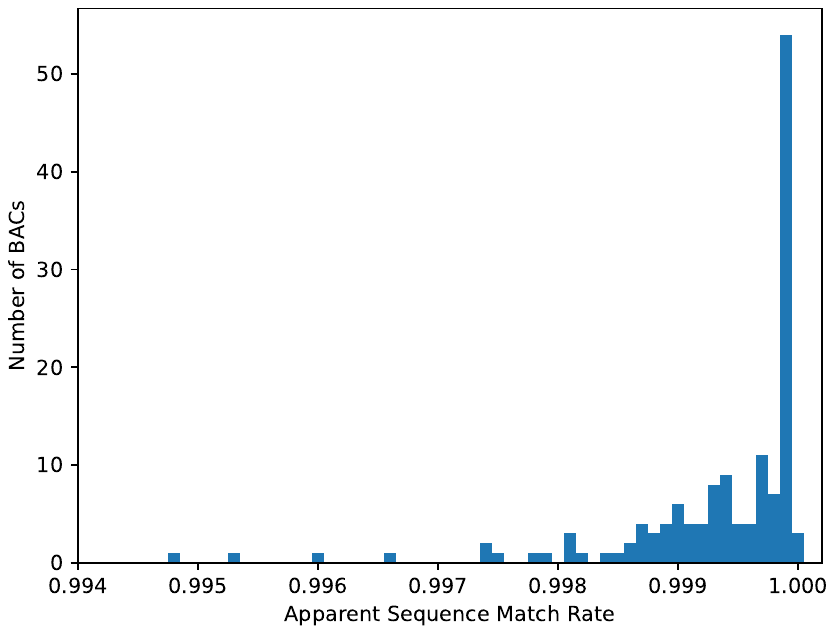

**Figure S3. Apparent sequence errors based on finished BAC clones**

The sequence of 142 finished BAC clones from the CH-82 library were aligned to the Dog_10k_Boxer_Tasha_1.0 assembly. Depicted is a histogram of the alignment identity (ignoring gaps) for each clone. The overall match rate is 0.9995, corresponding to a Phred-scaled quality value of Q33.

| a. | 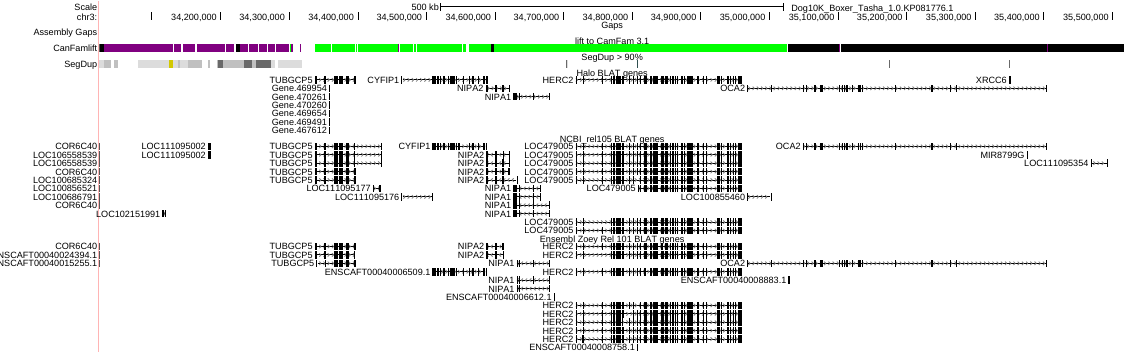 |
| --- | --- |
| b. | 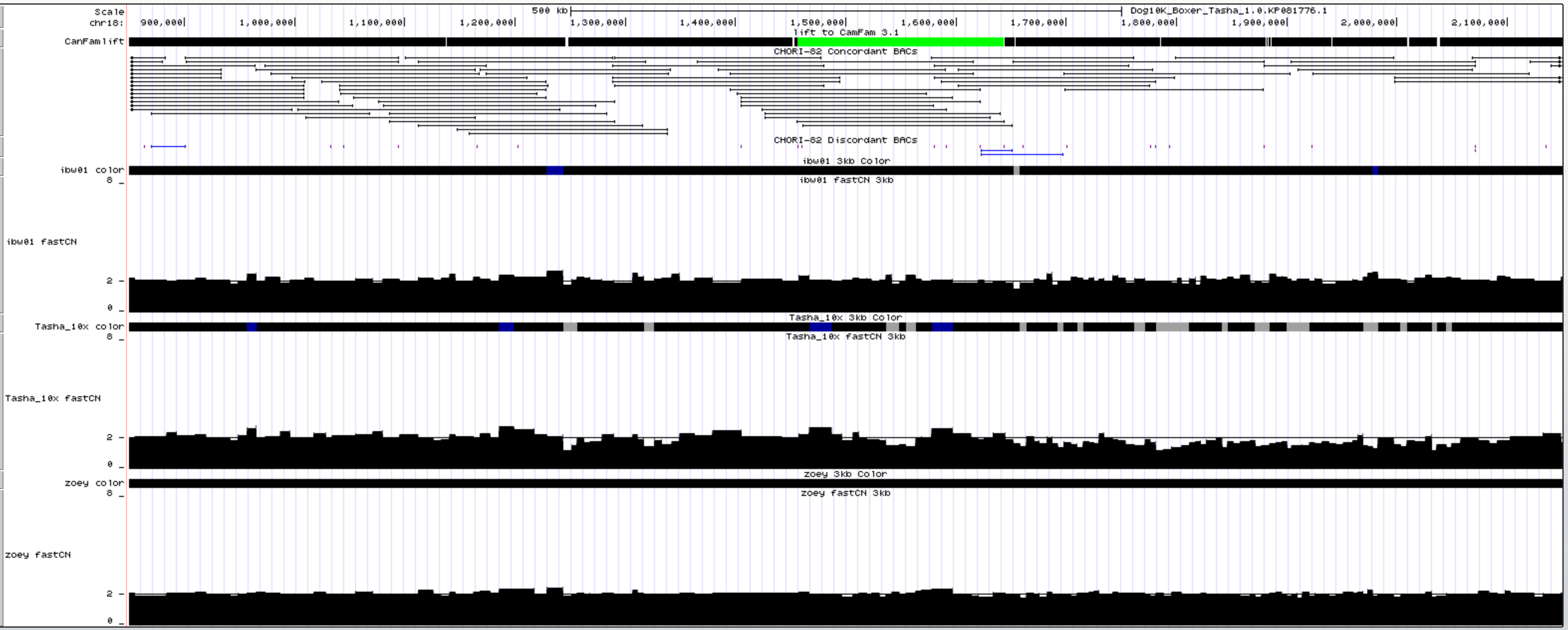 |
| c. | 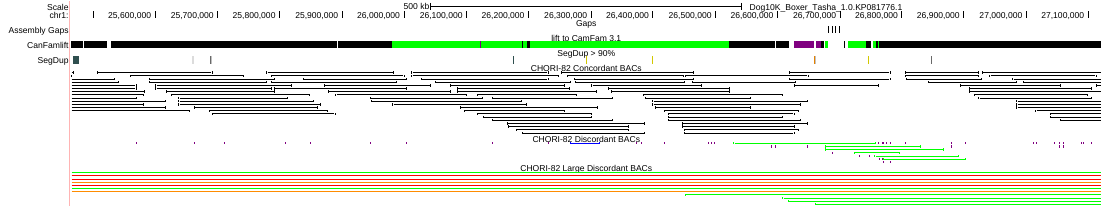 |

**Figure S4.** Correction of CanFam3.1 assembly errors documented by the correct alignment of CH-82 BAC end sequences. (**a**) *OCA2* gene (Chr3:34,350,000-35,050,000). The green line shows an incorrect inversion in CanFam3.1 compared to the new Dog_10k_Boxer_Tasha_1.0 assembly. (**b**) Chr18:850,000-2,150,000 region which was also misassembled in CanFam3.1 is now supported in the new assembly by concordantly aligned BAC-ends. (**c**) Chr1 region:25,463,622-27,120,619, another incorrect inversion in CanFam3.1, which is now correctly assembled in Dog10k_Boxer_Tasha_1.0.

| *COL9A3* | 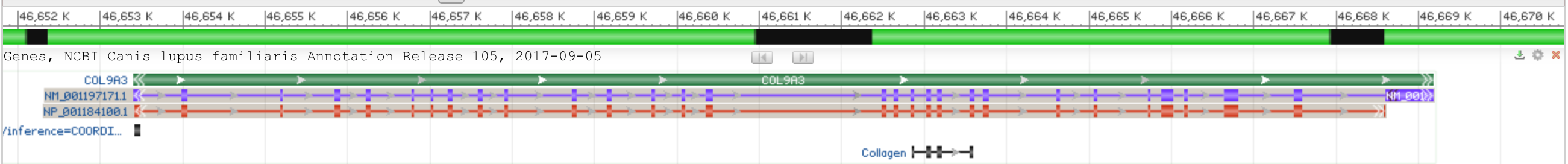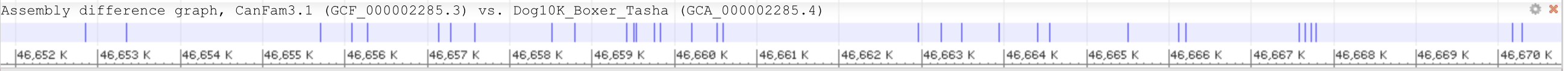 |
| --- | --- |
| *SCARF2* |  |
| *RASGRP1* | 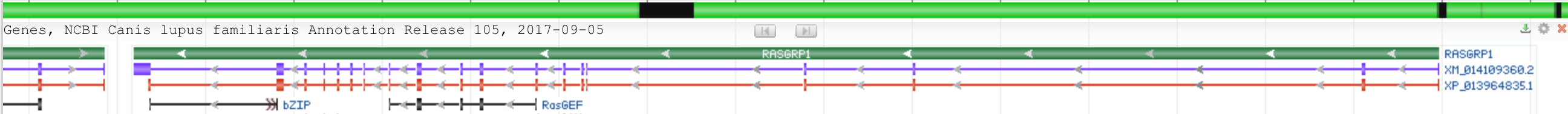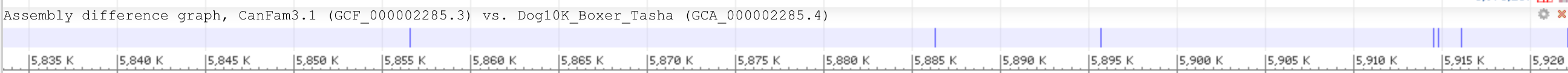 |
| *KLF4* | 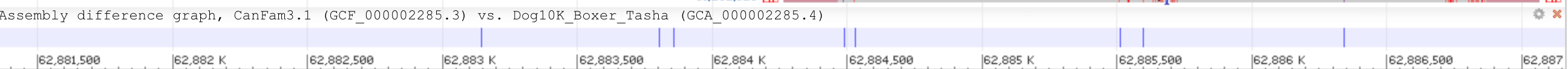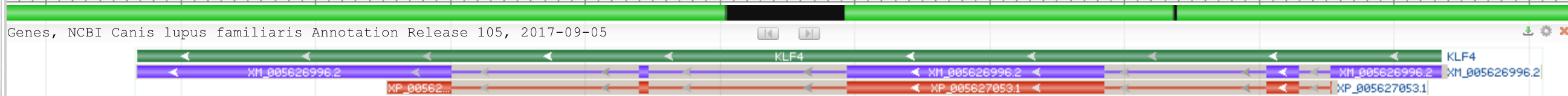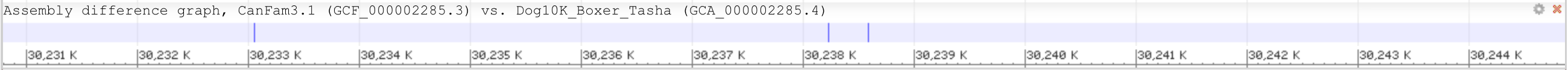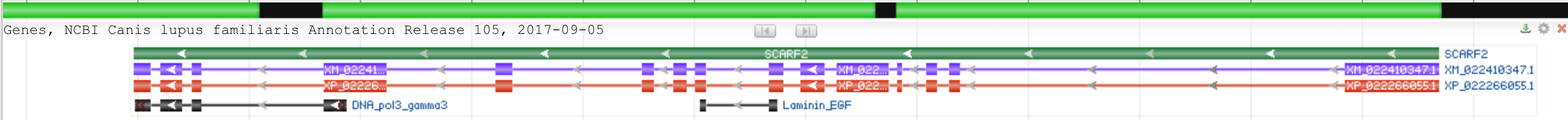 |

**Figure S5.** Gaps in exons of coding genes that have been filled in the Dog10k_Boxer_Tasha_1.0 assembly.

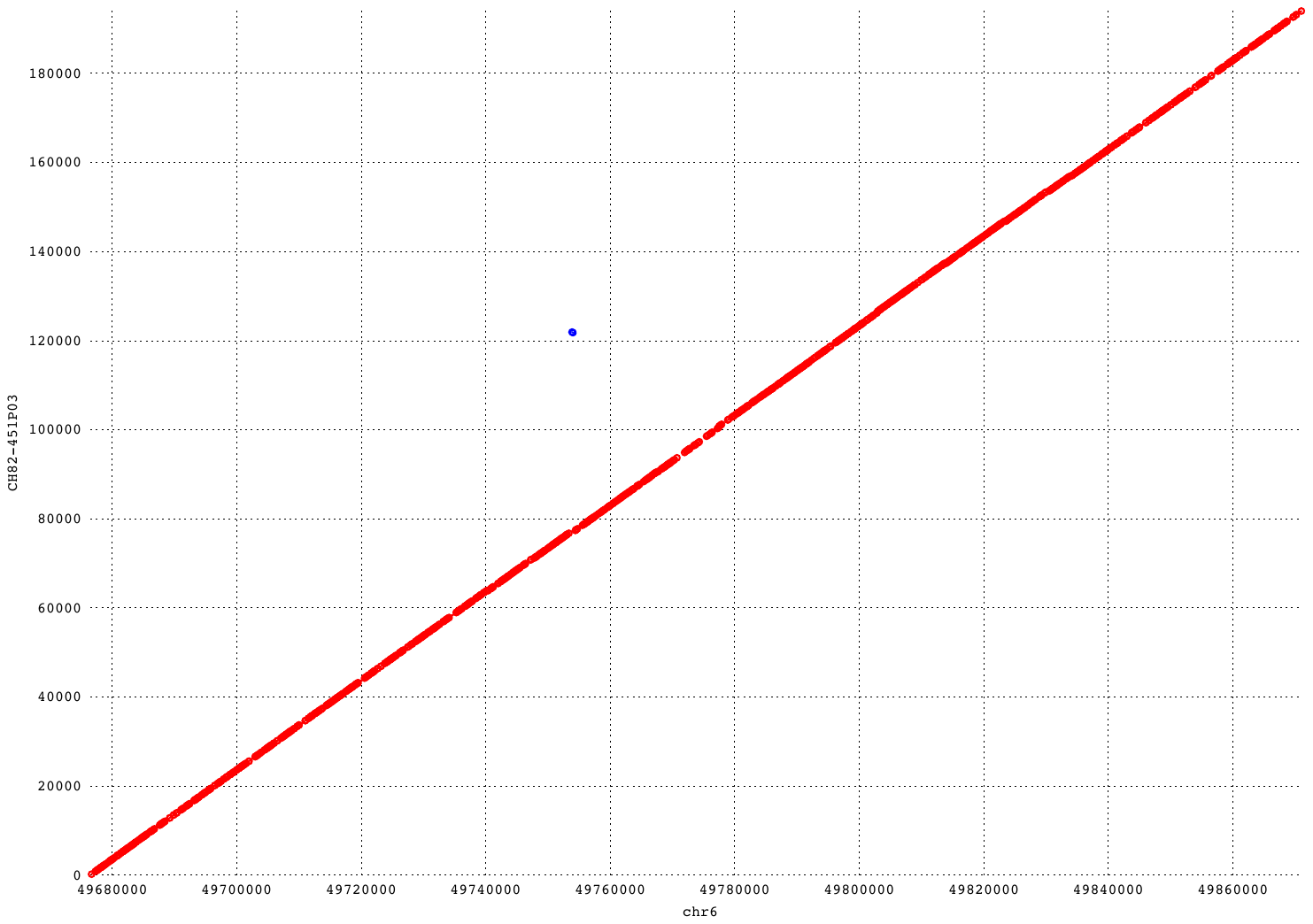

**Figure S6. Alignment of CH82-451P03 to the Dog10k_Boxer_Tasha_1.0 assembly**

The fully sequenced BAC clone was aligned to the chr6 from the Tasha4 assembly using mummer. Segments with a minimum match of 100 bp are depicted as red (forward orientation) or blue (reverse orientation) segments.
